# polars-bio – fast, scalable and out-of-core operations on large genomic interval datasets

**DOI:** 10.1101/2025.03.21.644629

**Authors:** Marek Wiewiórka, Pavel Khamutou, Marek Zbysiński, Tomasz Gambin

## Abstract

**Motivation:** Genomic studies very often rely on computational intensive analyses of relationships between features, which are typically represented as intervals along one-dimensional coordinate system (such as positions on a chromosome). In this context, Python programming language is extensively used for manipulation and analysis of data stored in a tabular-form of rows and columns called dataframe. Pandas is the most-widely used Python dataframe package and is criticized for its inefficiencies and scalability problems that its novel alternative – Polars – tries to address with a vectorized backend written in Rust programming language.

**Results:** *polars-bio* is a Python library that enables fast, parallel and out-of-core operations on large genomic intervals datasets. Its main components are implemented in Rust, using the Apache DataFusion query engine and Apache Arrow for efficient data representation. It is compatible with Polars and Pandas DataFrame formats. Single-thread benchmarking results confirm that for operations such as *count overlaps* is 38x and *coverage* is 15x faster than the state-of-the-art *bioframe* library. Our implementation of *overlap* operation consumes 90x less memory in streaming mode. Multi-thread benchmarks show good scalability and up to 282x speedup for *count overlaps* operation when executed with 8 CPU cores. To the best of our knowledge, *polars-bio* is the fastest single-node library for genomic interval dataframes in Python.

**Availability and implementation:** *polars-bio* is an open source Python package distributed under the Apache License available for all main platforms, such as Linux, macOS and Windows in the PyPI registry. The web page is https://biodatageeks.org/polars-bio/ and the source code is available on GitHub: https://github.com/biodatageeks/polars-bio

## Introduction

Operations on genomic intervals (ranges), such as finding intervals that overlap with or are closest to a given interval, are fundamental to many bioinformatic analyses, including the process of annotating genetic variants[12]. Dedicated efficient data structures and/or accompanying command line interface tools, such as NCList[1], AIList[3], BedTools[16], Bedtk[10], Coitrees[6] have been proposed. In addition, growing popularity of the Python programming language for data analysis resulted in the emergence of libraries, such as pybedtools[2], Python port of GenominRanges[9] or more recently PyRanges[17], Pygenomics[18] and Bioframe[14]. The existing approaches, however, suffer from one or more problems that include:

1. not adhering to standardized Python data formats, like Pandas/Polars DataFrames, and widely-adopted efficient data exchange methods, like Arrow C data interface, which results in costly in-memory data transfers,

- tool-specific data representation hinders seamless integration with the rich Python analytics ecosystem,
- suboptimal performance due to the use of inefficient data structures or memory representation and missing vectorized execution engine,
- lack or inefficient parallel and out-of-core computational models for large scale datasets,
- missing capability of streaming processing and federated queries of large-scale datasets directly from cloud storage services without the need of downloading them first. In many cases this approach might be not feasible due to the datasets size that can exceed both computer’s main memory but also locally attached drives capacity. Such a functionality is beneficial as many research projects share their datasets using services like Amazon Simple Storage Service (S3) or Google Cloud Storage (GCS).

To address the aforementioned challenges, we present *polars-bio*, a Python library that combines the strengths of Apache DataFusion[8] – extensible, columnar, streaming, multi-threaded, vectorized execution engine, Apache Arrow[5] – columnar memory format enabling efficient data representation and exchange coupled with the user-friendly and fast Polars[13] library.

## Methods

### Architecture

When designing *polars-bio* we took inspiration from the recently proposed paradigm of Composable Data Management Systems[15] and built it upon modular and reusable components to achieve standardization of **language fronted** (Structured Query Language (SQL) and DataFrame), extensibility of **query optimization** and high-performance of **execution engine**. From the data processing perspective, we adopted **lazy evaluation, asynchronous, streaming** and **parallel** programming patterns to enable large-scale computations and resources usage efficiency. *polars-bio* comprises the following elements:

- Internal data representation – Apache Arrow is used as a columnar memory model to facilitate efficient (in most of the cases zero-copy) data exchange between components in a form of streams of record batches (i.e. arrays of array references representing typed columns with Arrow C data/stream interfaces) (see Fig. 1A point **4a** and **4b**). It is feasible because both Pandas and Polars also use Arrow-compatible memory layouts (Fig. 1A - **3**).
- Parallel query engine (Fig. 1A - **8**) – Apache DataFusion was selected due to its great extensibility (e.g. user-defined functions/table functions, custom table providers, logical and physical plan optimization rules or execution plan operators), high efficiency (based on asynchronous and parallel execution model processing streams of partitioned dataframes represtented logically as tables backed by different providers – Fig. 1A - **3**) and finally active community of open-source contributors and companies that embedded it in their specialized data management systems.
- Native file formats readers and tables providers – Apache DataFusion has built-in support for popular file formats, such as comma-separated values (CSV), Parquet. For bioinformatic file formats, such as Variant Call Format for variants (VCF) or Browser Extensible Data (BED), custom tables providers based on the noodles[11] library have been developed. They are also implemented in an asynchronous streaming fashion – Fig. 1A - **2**.
- Query planner extensions – a set of custom physical optimization rules for interval operations has been added to detect and override default operators (based on hash or nested-loop join algorithms), so that our more efficient, interval trees-based methods are used instead – Fig. 1A - **6**.
- User-defined table functions – for range operations, such as *coverage* or *count overlaps*, that cannot be efficiently expressed with either DataFusion DataFrame or SQL application programming interfaces (APIs) – Fig. 1A - **7**.
- Data access layer – in order to provide a unified access to network storage systems (public cloud object storages, e.g. S3 or GCS) Apache DAL project was used as an abstraction layer of data streams over the existing cloud-specific access methods – Fig. 1A - **1**.
- Language fronted – by adhering to the popular Polars and Pandas APIs, users can easily plug in *polars-bio* to the existing bioinformatic analyses with few code changes required. Besides, SQL API can be used for efficient pre-processing of input files.

**Fig. 1.**
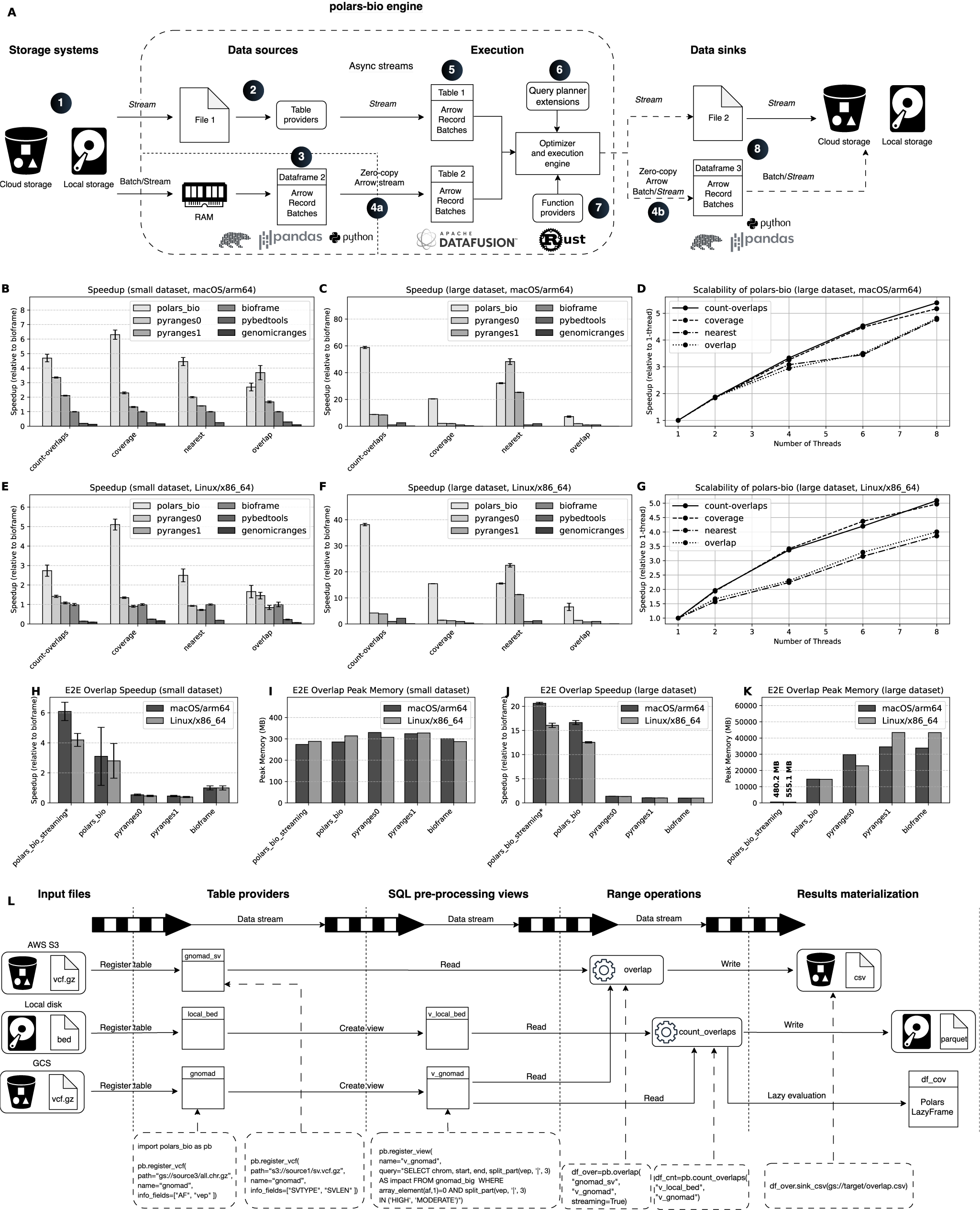
Panel A shows main components of *polars-bio* architecture (see Architecture section for details). Panels B–G show performance benchmarks: Panels B, C, E, and F display speedup (relative to bioframe, single thread) on small and large datasets for macOS/arm64 and Linux/x86 64, while Panels D and G present scalability results for *polars-bio* (scalability was evaluated as speedup versus the number of threads). Panels H–K show performance results for end-to-end processing of overlap operation: Panels H and J present speedup, while panels I and K illustrate peak memory usage, * - in *polars bio streaming* despite limiting the number of threads in DataFusion and in Polars, CPU utilization was constant and approximately 160%. Panel L shows an example of a practical use case scenario (see Functionality section for details).

## Implementation

*polars-bio* is implemented as Python package with a Rust backend that is distributed in a form of platform-specific binaries targeted for Linux (x86 64), macOS (x86 64 and arm64) and Windows (x86 64) platforms. It is developed using PyO3[4] Rust bindings for Python.

## Functionality

### Supported Operations and Input formats

*polars-bio* provides a rich set of genomics operations. *overlap* determines which genomic intervals from one dataset overlap those in another one. *nearest* takes two sets of intervals (say A and B) and for each interval *a* in A finds the interval in B that either overlaps with *a* or is closest to *a*. It could be used to find the nearest gene to a genetic variant. *count overlaps* calculates, for each interval in a primary set A, how many intervals from a secondary set B overlap it. It is useful for applications such as counting how many reads (intervals in B) cover each gene (interval in A), or how many regulatory elements fall within each genomic bin. *coverage* calculates the number of base pairs of A covered by intervals in B. See online documentation for all supported operations.

To facilitate real-world genomics workflows, *polars-bio* includes built-in readers for standard genomic file formats, such as BED (for genomic intervals), VCF, BAM (alignment reads) and FASTQ/FASTA (sequence reads and reference sequences). *polars-bio* supports both 0-based and 1-based coordinate systems, allowing users to specify the desired system as a parameter for each operation.

### Practical usecases

*polars-bio* exposes an API that works with both Pandas and Polars. Users can call high-level functions (e.g. *overlap* or *count overlaps*) or use its SQL interface (powered by Apache DataFusion) to register datasets as in-memory table references. A major strength is its support for federated analysis, processing data from multiple sources without central consolidation.

For example, consider: (i) a local BED file of genomic intervals; (ii) 1M structural variants (~2 GB) from gnomAD[7] stored on GCS; and 900M SNVs and indels (~500 GB) on Amazon S3 (see Fig.1L).

Traditionally, one would need to download these files or use a distributed query engine. With *polars-bio*, one can register the remote files as tables and: (i) parse specific columns (e.g. extract the IMPACT value from Variant Effect Predictor annotations); (ii) filter variants via SQL to select those that are rare and have HIGH or MODERATE impact; (iii) overlap the filtered variants with structural variants. Reading, preprocessing and overlap operations are executed in streaming mode: *polars-bio* reads chunks from the remote files, transforms and filters them, finds overlaps and writes results (to CSV or Parquet) without ever loading the full 500 GB into memory. One can also easily count overlaps between a set of genomic intervals defined in local BED file and remote variant data to determine the number of variants in each genomic region.

## Performance

*polars-bio* has been carefully benchmarked for speed, scalability and memory consumption on two popular platforms: Linux with x86 64 and macOS with arm64 architectures. The benchnmarking datasets proposed in the AIList manuscript has been used in all the tests (the “small” test-case used datasets 1,2 whereas the “large” one – 8,7). In a single-threaded mode the following Python libraries have been compared: bioframe, pyranges (major version 0 and 1 that is complete rewrite of the former one), pybedtools and GenomicRanges running 4 popular operations: *overlap, nearest, count overlaps* and *coverage*. In 3 out of 4 tests polars-bio proved to be the fastest tool on both small (with average speedup of all operations versus bioframe: vary from ~3.0x to ~4.5x depending on computer architecture) and large (average speedup: from ~18.9x to ~29.6x) datasets (Fig. 1B,C,E,F). Additionally, an end-to-end test scenario that encompassed reading from Parquet files, computing interval overlaps and saving results to CSV files was profiled (Fig. 1H- K). In this benchmark *polars-bio* has been run twice: with full results materialization (polars bio) in RAM prior saving the output and in streaming mode (polars bio streaming) when only current subset of record batches were materialized. In these tests both modes (full materialization and streaming) *polars-bio* were significantly faster than bioframe and pyranges. For small dataset *polars-bio* achieved speedups from ~2.8x to ~6.1x, while for large dataset speedup varied between ~12.6x and ~20.6x (Fig. 1H,J). For a small test scenario memory usage was comparable across all tools, in the case of large dataset one can observe clear advantage of *polars-bio*: in streaming mode memory utilization on macOS was measured ~560MB whereas bioframe at peak needed ~60x, pyranges ~54x and pyranges1 ~62x more. In the case of Linux, the factors were: ~90x, ~48x and ~90x, respectively. In both benchmarks *polars-bio* in the default mode required 14.6-14.7 GB of RAM at peak (Fig. 1I,K). See supplementary materials for complete benchmarking results, testing environment software and hardware details.

## Conclusion

The *polars-bio* bridges the gap between the efficiency of command-line tools written in natively compiled languages, such as C++ or Rust, and ergonomics of the Python dataframes ecosystem. Out-of-core (streaming) processing feature addresses in many cases the constantly growing need for Big Data and federated queries capabilities without incurring the complexity of the distributed computing machinery. It enables both *adhoc* exploratory analyses and recurrent data pipelines for small and large datasets to be implemented in a fast, scalable and consistent way.

