## supplementary materials for "polars-bio – fast, scalable and out-of-core operations on large genomic interval datasets"

### 🔬 Supplementary material

#### Supplemental material

This document provides additional information about the benchmarking setup, data, and results that were presented in the manuscript.

#### Benchmark setup

##### Code and benchmarking scenarios

Repository

##### Operating systems and hardware configurations

**macOS**

- cpu architecture: `arm64`
- cpu name: `Apple M3 Max`
- cpu cores: `16`
- memory: `64 GB`
- kernel: `Darwin Kernel Version 24.2.0: Fri Dec  6 19:02:12 PST 2024; root:xnu-11215.61.5~2/RELEASE_ARM64_T6031`
- system: `Darwin`
- os-release: `macOS-15.2-arm64-arm-64bit`
- python: `3.12.4`
- polars-bio: `0.8.3`

**Linux**

c3-standard-22 machine was used for benchmarking.

- cpu architecture: `x86_64`
- cpu name: `Intel(R) Xeon(R) Platinum 8481C CPU @ 2.70GHz`

- cpu cores: `22`
- memory: `88 GB`
- kernel: `Linux-6.8.0-1025-gcp-x86_64-with-glibc2.35`
- system: `Linux`
- os-release: `#27~22.04.1-Ubuntu SMP Mon Feb 24 16:42:24 UTC 2025`
- python: `3.12.8`
- polars-bio: `0.8.3`

#### Software

- Bioframe-0.7.2
- PyRanges0-0.0.132
- PyRanges1-e634a11
- pybedtools-0.10.0
- PyGenomics-0.1.1
- GenomicRanges-0.5.0

#### Data

AIList dataset was used for benchmarking.

| Dataset# | Name | Size(x1000) | Non-flatness |
|---|---|---|---|
| 0 | chainRn4 | 2,351 | 6 |
| 1 | fBrain | 199 | 1 |
| 2 | exons | 439 | 2 |
| 3 | chainOrnAna1 | 1,957 | 6 |
| 4 | chainVicPac2 | 7,684 | 8 |
| 5 | chainXenTro3Link | 50,981 | 7 |

| | | | |
|---|---|---|---|
| 7 | ex-anno | 1,194 | 2 |
| 8 | ex-rna | 9,945 | 7 |

**Note**

Test dataset in *Parquet* format can be downloaded from:

- databio.zip

#### Single thread results

Results for `overlap`, `nearest`, `count-overlaps`, and `coverage` operations with single-thread performance on `apple-m3-max` and `gcp-linux` platforms.

###### apple-m3-max

#### 1-2

OVERLAP

| Library | Min (s) | Max (s) | Mean (s) | Speedup |
|---|---|---|---|---|
| polars_bio | 0.035619 | 0.043113 | 0.0383 | 2.70x |
| bioframe | 0.102257 | 0.104425 | 0.103354 | 1.00x |
| pyranges0 | 0.025425 | 0.032821 | 0.028001 | 3.69x |
| pyranges1 | 0.059608 | 0.064147 | 0.061763 | 1.67x |
| pybedtools | 0.343204 | 0.352804 | 0.348434 | 0.30x |
| genomicranges | 1.042893 | 1.044245 | 1.043488 | 0.10x |

NEAREST

| Library | Min (s) | Max (s) | Mean (s) | Speedup |
|---|---|---|---|---|

| | | | | |
|---|---|---|---|---|
| polars_bio | 0.039943 | 0.045166 | 0.042109 | 4.45x |
| bioframe | 0.185452 | 0.189631 | 0.187388 | 1.00x |
| pyranges0 | 0.092334 | 0.09634 | 0.093688 | 2.00x |
| pyranges1 | 0.133631 | 0.134179 | 0.133981 | 1.40x |
| pybedtools | 0.756676 | 0.761866 | 0.75953 | 0.25x |

###### COUNT-OVERLAPS

| Library | Min (s) | Max (s) | Mean (s) | Speedup |
|---|---|---|---|---|
| polars_bio | 0.026706 | 0.029754 | 0.028142 | 4.69x |
| bioframe | 0.131124 | 0.133729 | 0.132052 | 1.00x |
| pyranges0 | 0.039136 | 0.039774 | 0.039377 | 3.35x |
| pyranges1 | 0.061976 | 0.063181 | 0.062658 | 2.11x |
| pybedtools | 0.665804 | 0.673844 | 0.668534 | 0.20x |
| genomicranges | 0.994963 | 1.006435 | 0.999389 | 0.13x |

###### COVERAGE

| Library | Min (s) | Max (s) | Mean (s) | Speedup |
|---|---|---|---|---|
| polars_bio | 0.0262 | 0.028749 | 0.027418 | 6.30x |
| bioframe | 0.16949 | 0.176628 | 0.172842 | 1.00x |
| pyranges0 | 0.07376 | 0.076708 | 0.075369 | 2.29x |
| pyranges1 | 0.128027 | 0.133263 | 0.130247 | 1.33x |

| | | | | |
|---|---|---|---|---|
| pybedtools | 0.701817 | 0.708726 | 0.705839 | 0.24x |
| genomicranges | 1.032651 | 1.049059 | 1.040799 | 0.17x |

### 8-7

###### OVERLAP

| Library | Min (s) | Max (s) | Mean (s) | Speedup |
|---|---|---|---|---|
| polars_bio | 3.987391 | 4.648581 | 4.235518 | 7.17x |
| bioframe | 29.793837 | 30.991576 | 30.375518 | 1.00x |
| pyranges0 | 15.632212 | 15.974075 | 15.857213 | 1.92x |
| pyranges1 | 31.622804 | 33.699074 | 32.680701 | 0.93x |
| pybedtools | 916.711575 | 919.974811 | 918.154834 | 0.03x |
| genomicranges | 479.214112 | 487.832054 | 484.579554 | 0.06x |

###### NEAREST

| Library | Min (s) | Max (s) | Mean (s) | Speedup |
|---|---|---|---|---|
| polars_bio | 2.116922 | 2.169534 | 2.139006 | 32.13x |
| bioframe | 68.581465 | 68.992651 | 68.725495 | 1.00x |
| pyranges0 | 1.381964 | 1.508513 | 1.424446 | 48.25x |
| pyranges1 | 2.697684 | 2.728407 | 2.717532 | 25.29x |
| pybedtools | 35.528719 | 35.876667 | 35.699544 | 1.93x |

###### COUNT-OVERLAPS

| Library | Min (s) | Max (s) | Mean (s) | Speedup |
|---|---|---|---|---|
| polars_bio | 1.445467 | 1.484052 | 1.46225 | 58.77x |
| bioframe | 85.632767 | 86.26148 | 85.935955 | 1.00x |
| pyranges0 | 9.674847 | 9.833233 | 9.753982 | 8.81x |
| pyranges1 | 10.170249 | 10.254359 | 10.201813 | 8.42x |
| pybedtools | 33.101592 | 33.966188 | 33.423595 | 2.57x |
| genomicranges | 488.972732 | 490.395787 | 489.548184 | 0.18x |

COVERAGE

| Library | Min (s) | Max (s) | Mean (s) | Speedup |
|---|---|---|---|---|
| polars_bio | 1.195279 | 1.205765 | 1.199323 | 20.45x |
| bioframe | 24.423391 | 24.682901 | 24.525909 | 1.00x |
| pyranges0 | 11.093644 | 11.328071 | 11.220416 | 2.19x |
| pyranges1 | 11.987003 | 12.147925 | 12.066045 | 2.03x |
| pybedtools | 59.699275 | 60.04087 | 59.84965 | 0.41x |
| genomicranges | 500.041974 | 503.31936 | 502.043072 | 0.05x |

##### gcp-linux

### 1-2

OVERLAP

| Library | Min (s) | Max (s) | Mean (s) | Speedup |
|---|---|---|---|---|

| | | | | |
|---|---|---|---|---|
| polars_bio | 0.045943 | 0.064732 | 0.054234 | 1.66x |
| bioframe | 0.084137 | 0.099481 | 0.090107 | 1.00x |
| pyranges0 | 0.056206 | 0.065654 | 0.061844 | 1.46x |
| pyranges1 | 0.09908 | 0.119018 | 0.106228 | 0.85x |
| pybedtools | 0.38246 | 0.406379 | 0.39153 | 0.23x |
| genomicranges | 1.19939 | 1.224621 | 1.208255 | 0.07x |

NEAREST

| Library | Min (s) | Max (s) | Mean (s) | Speedup |
|---|---|---|---|---|
| polars_bio | 0.057012 | 0.073822 | 0.064665 | 2.49x |
| bioframe | 0.158764 | 0.165707 | 0.161273 | 1.00x |
| pyranges0 | 0.172297 | 0.176259 | 0.17363 | 0.93x |
| pyranges1 | 0.217619 | 0.234088 | 0.22335 | 0.72x |
| pybedtools | 0.845945 | 0.84898 | 0.847447 | 0.19x |

COUNT-OVERLAPS

| Library | Min (s) | Max (s) | Mean (s) | Speedup |
|---|---|---|---|---|
| polars_bio | 0.035631 | 0.043555 | 0.04066 | 2.74x |
| bioframe | 0.108015 | 0.116522 | 0.111266 | 1.00x |
| pyranges0 | 0.077336 | 0.080282 | 0.07844 | 1.42x |
| pyranges1 | 0.100883 | 0.106671 | 0.103181 | 1.08x |

| | | | | |
|---|---|---|---|---|
| pybedtools | 0.745958 | 0.759006 | 0.754393 | 0.15x |
| genomicranges | 1.154942 | 1.164158 | 1.158506 | 0.10x |

COVERAGE

| Library | Min (s) | Max (s) | Mean (s) | Speedup |
|---|---|---|---|---|
| polars_bio | 0.036476 | 0.040001 | 0.037897 | 5.10x |
| bioframe | 0.189201 | 0.20046 | 0.193401 | 1.00x |
| pyranges0 | 0.141659 | 0.14424 | 0.143188 | 1.35x |
| pyranges1 | 0.206033 | 0.224902 | 0.213089 | 0.91x |
| pybedtools | 0.773732 | 0.780424 | 0.776934 | 0.25x |
| genomicranges | 1.186341 | 1.194172 | 1.189255 | 0.16x |

### 8-7

OVERLAP

| Library | Min (s) | Max (s) | Mean (s) | Speedup |
|---|---|---|---|---|
| polars_bio | 6.235223 | 9.61441 | 7.723144 | 6.54x |
| bioframe | 50.319263 | 50.956633 | 50.537202 | 1.00x |
| pyranges0 | 36.371926 | 36.581642 | 36.448645 | 1.39x |
| pyranges1 | 63.336711 | 63.455435 | 63.40654 | 0.80x |
| pybedtools | 1149.001487 | 1152.127068 | 1150.070659 | 0.04x |
| genomicranges | 597.951648 | 599.960895 | 599.002871 | 0.08x |

NEAREST

| Library | Min (s) | Max (s) | Mean (s) | Speedup |
|---|---|---|---|---|
| polars_bio | 3.576373 | 3.679698 | 3.633697 | 15.54x |
| bioframe | 56.301865 | 56.776617 | 56.464305 | 1.00x |
| pyranges0 | 2.45308 | 2.60494 | 2.505172 | 22.54x |
| pyranges1 | 4.975662 | 5.011008 | 4.997007 | 11.30x |
| pybedtools | 44.181913 | 44.79409 | 44.386971 | 1.27x |

COUNT-OVERLAPS

| Library | Min (s) | Max (s) | Mean (s) | Speedup |
|---|---|---|---|---|
| polars_bio | 2.052196 | 2.104447 | 2.075706 | 38.15x |
| bioframe | 79.174164 | 79.234115 | 79.194209 | 1.00x |
| pyranges0 | 18.797436 | 18.851941 | 18.824498 | 4.21x |
| pyranges1 | 20.399172 | 20.436149 | 20.418562 | 3.88x |
| pybedtools | 35.850631 | 36.142479 | 36.041115 | 2.20x |
| genomicranges | 612.985873 | 613.52087 | 613.229997 | 0.13x |

COVERAGE

| Library | Min (s) | Max (s) | Mean (s) | Speedup |
|---|---|---|---|---|
| polars_bio | 1.829478 | 1.838981 | 1.834999 | 15.44x |
| bioframe | 28.29136 | 28.361417 | 28.326821 | 1.00x |

| | | | | |
|---|---|---|---|---|
| pyranges0 | 18.611247 | 20.021441 | 19.473105 | 1.45x |
| pyranges1 | 22.118838 | 22.210733 | 22.161329 | 1.28x |
| pybedtools | 74.477086 | 74.868659 | 74.618066 | 0.38x |
| genomicranges | 623.865655 | 623.94955 | 623.896645 | 0.05x |

#### Parallel performance

Results for parallel operations with 1, 2, 4, 6 and 8 threads.

**apple-m3-max**

**8-7**

OVERLAP

| Library | Min (s) | Max (s) | Mean (s) | Speedup |
|---|---|---|---|---|
| polars_bio | 3.247022 | 3.803021 | 3.370889 | 1.00x |
| polars_bio-2 | 1.798569 | 1.848162 | 1.811417 | 1.86x |
| polars_bio-4 | 1.140229 | 1.158243 | 1.147355 | 2.94x |
| polars_bio-6 | 0.959703 | 0.968725 | 0.962915 | 3.50x |
| polars_bio-8 | 0.694637 | 0.710492 | 0.701048 | 4.81x |

NEAREST

| Library | Min (s) | Max (s) | Mean (s) | Speedup |
|---|---|---|---|---|
| polars_bio | 2.186354 | 2.248171 | 2.220822 | 1.00x |
| polars_bio-2 | 1.162969 | 1.222115 | 1.187505 | 1.87x |

| | | | | |
|---|---|---|---|---|
| polars_bio-6 | 0.632877 | 0.652955 | 0.642816 | 3.45x |
| polars_bio-8 | 0.456674 | 0.476473 | 0.465284 | 4.77x |

COUNT-OVERLAPS

| Library | Min (s) | Max (s) | Mean (s) | Speedup |
|---|---|---|---|---|
| polars_bio | 1.502551 | 1.534006 | 1.515078 | 1.00x |
| polars_bio-2 | 0.811236 | 0.821365 | 0.815682 | 1.86x |
| polars_bio-4 | 0.440628 | 0.46778 | 0.455358 | 3.33x |
| polars_bio-6 | 0.331317 | 0.338207 | 0.334638 | 4.53x |
| polars_bio-8 | 0.280465 | 0.282707 | 0.281311 | 5.39x |

COVERAGE

| Library | Min (s) | Max (s) | Mean (s) | Speedup |
|---|---|---|---|---|
| polars_bio | 1.181806 | 1.185549 | 1.183889 | 1.00x |
| polars_bio-2 | 0.644288 | 0.645076 | 0.644587 | 1.84x |
| polars_bio-4 | 0.362752 | 0.363411 | 0.363036 | 3.26x |
| polars_bio-6 | 0.258583 | 0.272702 | 0.264111 | 4.48x |
| polars_bio-8 | 0.222888 | 0.234884 | 0.229052 | 5.17x |

##### gcp-linux

### 8-7

OVERLAP

| Library | Min (s) | Max (s) | Mean (s) | Speedup |
|---|---|---|---|---|
| polars_bio | 6.325617 | 8.185275 | 7.005925 | 1.00x |
| polars_bio-2 | 3.920645 | 4.617084 | 4.198055 | 1.67x |
| polars_bio-4 | 3.036273 | 3.060781 | 3.0452 | 2.30x |
| polars_bio-6 | 2.127994 | 2.134505 | 2.131016 | 3.29x |
| polars_bio-8 | 1.731485 | 1.789347 | 1.752986 | 4.00x |

NEAREST

| Library | Min (s) | Max (s) | Mean (s) | Speedup |
|---|---|---|---|---|
| polars_bio | 4.047329 | 4.439016 | 4.198198 | 1.00x |
| polars_bio-2 | 2.624132 | 2.722843 | 2.682361 | 1.57x |
| polars_bio-4 | 1.809028 | 1.917798 | 1.871763 | 2.24x |
| polars_bio-6 | 1.309557 | 1.362131 | 1.333989 | 3.15x |
| polars_bio-8 | 1.066945 | 1.113168 | 1.087907 | 3.86x |

COUNT-OVERLAPS

| Library | Min (s) | Max (s) | Mean (s) | Speedup |
|---|---|---|---|---|
| polars_bio | 2.426441 | 2.456318 | 2.439266 | 1.00x |
| polars_bio-2 | 1.22516 | 1.272066 | 1.245401 | 1.96x |
| polars_bio-4 | 0.711421 | 0.744023 | 0.724315 | 3.37x |
| polars_bio-6 | 0.563797 | 0.607321 | 0.580574 | 4.20x |

| | | | | |
|---|---|---|---|---|
| polars_bio-8 | 0.459308 | 0.493886 | 0.479126 | 5.09x |

COVERAGE

| Library | Min (s) | Max (s) | Mean (s) | Speedup |
|---|---|---|---|---|
| polars_bio | 2.212958 | 2.23035 | 2.222531 | 1.00x |
| polars_bio-2 | 1.132056 | 1.15405 | 1.146413 | 1.94x |
| polars_bio-4 | 0.645737 | 0.661564 | 0.652277 | 3.41x |
| polars_bio-6 | 0.50589 | 0.511256 | 0.50839 | 4.37x |
| polars_bio-8 | 0.439503 | 0.450924 | 0.447075 | 4.97x |

#### End to end tests

Results for an end-to-end test with calculating overlaps and saving results to a CSV file.

#### apple-m3-max

### 1-2

E2E-OVERLAP-CSV

| Library | Min (s) | Max (s) | Mean (s) | Speedup | Peak memory (MB) |
|---|---|---|---|---|---|
| polars_bio | 0.042378 | 0.130957 | 0.071929 | 3.10x | 285.468 |
| polars_bio_streaming | 0.035498 | 0.037438 | 0.036653 | 6.09x | 274.093 |
| bioframe | 0.208548 | 0.251457 | 0.223219 | 1.00x | 300.75 |
| pyranges0 | 0.409707 | 0.415361 | 0.412135 | 0.54x | 329.968 |

| | | | | | |
|---|---|---|---|---|---|
| pyranges1 | 0.47518 | 0.491508 | 0.482739 | 0.46x | 324.468 |

**8-7**

E2E-OVERLAP-CSV

| Library | Min (s) | Max (s) | Mean (s) | Speedup | Peak memory (MB) |
|---|---|---|---|---|---|
| polars_bio | 22.781745 | 23.916568 | 23.161559 | 16.64x | 14677.0468 |
| polars_bio_streaming | 18.501279 | 18.797602 | 18.676707 | 20.63x | 555.109 |
| bioframe | 383.108514 | 387.500069 | 385.309331 | 1.00x | 33806.062 |
| pyranges0 | 276.421312 | 279.839508 | 277.845198 | 1.39x | 29777.312 |
| pyranges1 | 355.703878 | 367.680249 | 360.875151 | 1.07x | 34526.859 |

#### gcp-linux

**1-2**

E2E-OVERLAP-CSV

| Library | Min (s) | Max (s) | Mean (s) | Speedup | Peak memory (MB) |
|---|---|---|---|---|---|
| polars_bio | 0.072393 | 0.151871 | 0.09916 | 2.80x | 314.234 |
| polars_bio_streaming | 0.064092 | 0.067914 | 0.066202 | 4.19x | 288.621 |
| bioframe | 0.258278 | 0.31288 | 0.277225 | 1.00x | 287.101 |
| pyranges0 | 0.591745 | 0.599954 | 0.595204 | 0.47x | 307.218 |

### 8-7

E2E-OVERLAP-CSV

| Library | Min (s) | Max (s) | Mean (s) | Speedup | Peak memory (MB) |
|---|---|---|---|---|---|
| polars_bio | 44.539766 | 45.543038 | 45.196903 | 12.55x | 14575.14 |
| polars_bio_streaming | 34.007093 | 35.972075 | 35.309756 | 16.06x | 480.207 |
| bioframe | 566.167037 | 567.617695 | 567.13069 | 1.00x | 43295.378 |
| pyranges0 | 417.291061 | 421.875539 | 419.571591 | 1.35x | 22915.917 |
| pyranges1 | 538.365637 | 548.624613 | 543.918168 | 1.04x | 43408.699 |

###### Memory profiles

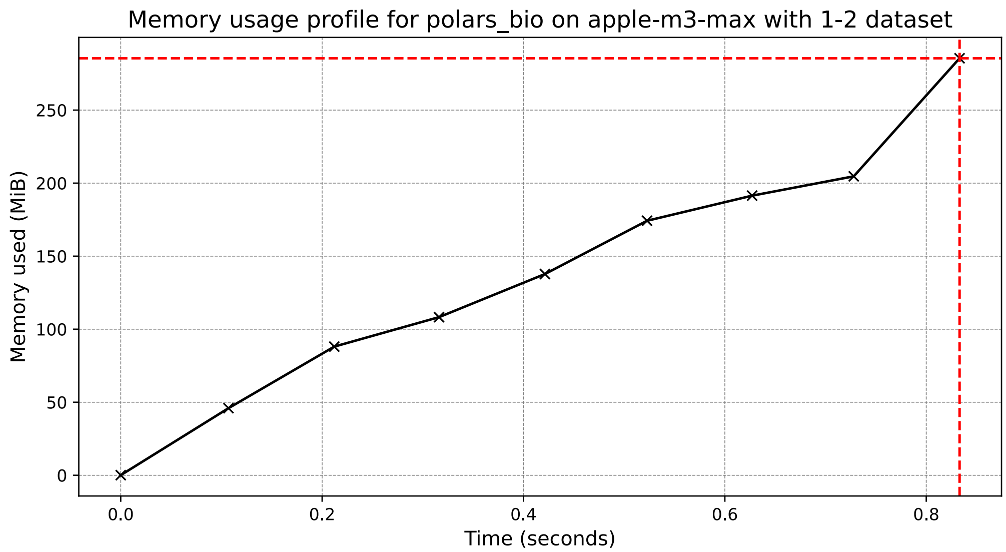
Memory usage profile for polars_bio on apple-m3-max with 1-2 dataset
Memory used (MiB)
0
50
100
150
200
250
0.0
0.2
0.4
0.6
0.8
Time (seconds)

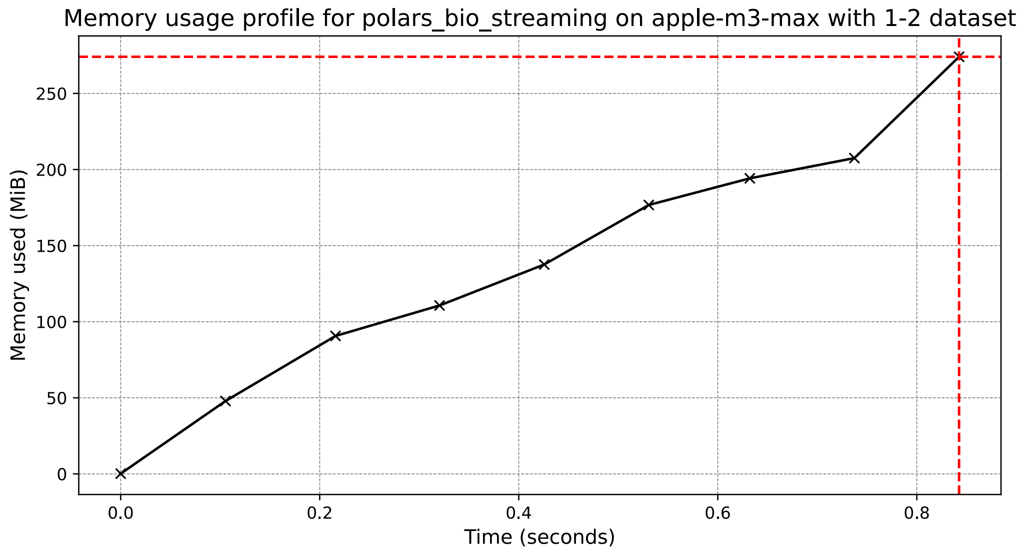
Memory usage profile for polars_bio_streaming on apple-m3-max with 1-2 dataset
Memory used (MiB)
0
50
100
150
200
250
0.0
0.2
0.4
0.6
0.8
Time (seconds)

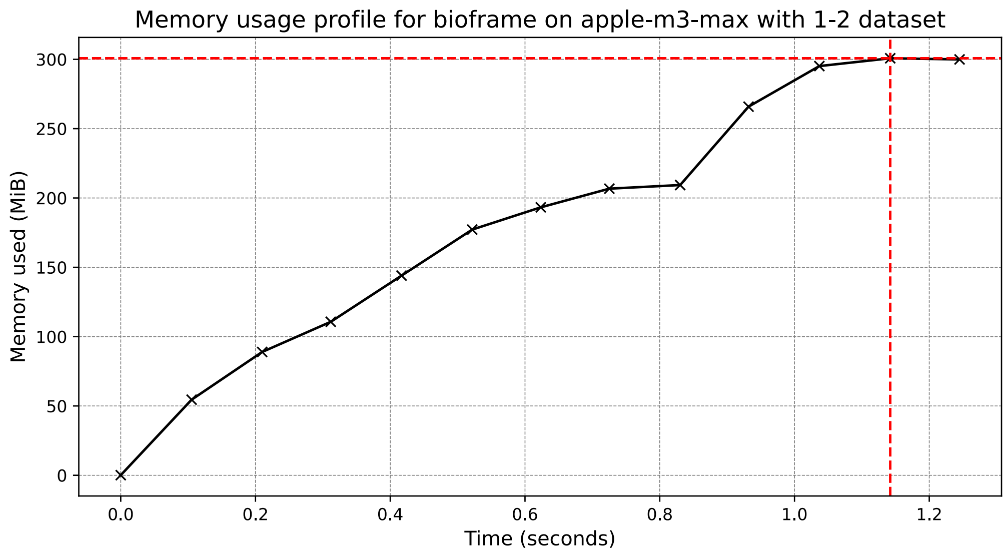

Memory usage profile for bioframe on apple-m3-max with 1-2 dataset
Memory used (MiB)
Time (seconds)
0
50
100
150
200
250
300
0.0
0.2
0.4
0.6
0.8
1.0
1.2


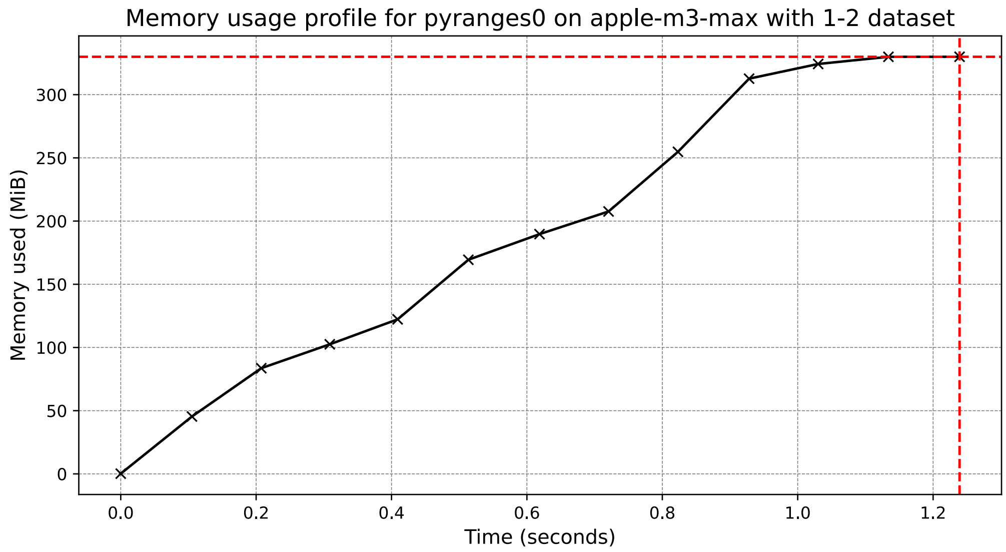

Memory usage profile for pyranges0 on apple-m3-max with 1-2 dataset
Memory used (MiB)
Time (seconds)
0
50
100
150
200
250
300
0.0
0.2
0.4
0.6
0.8
1.0
1.2

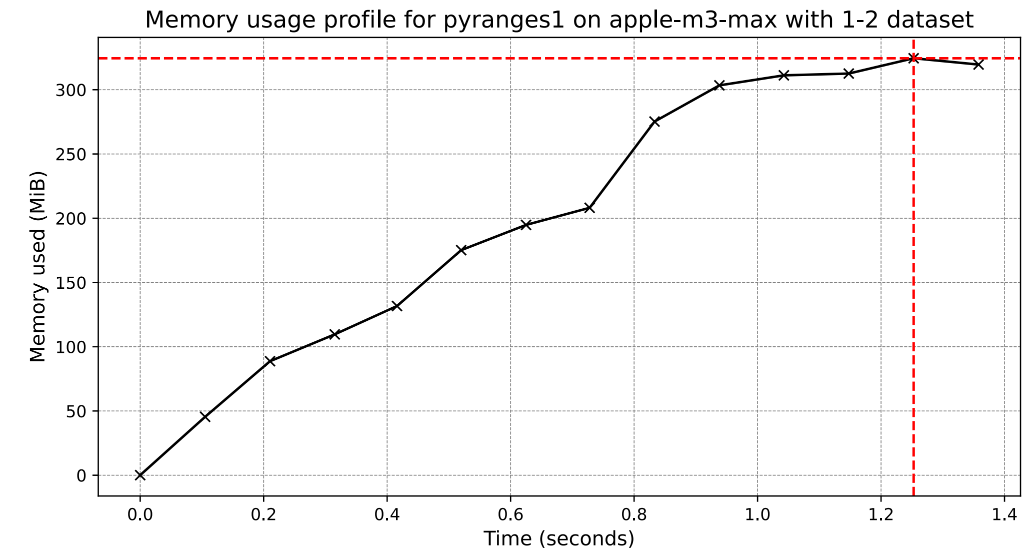
Memory usage profile for pyranges1 on apple-m3-max with 1-2 dataset
Memory used (MiB)
Time (seconds)
0
50
100
150
200
250
300
0.0
0.2
0.4
0.6
0.8
1.0
1.2
1.4

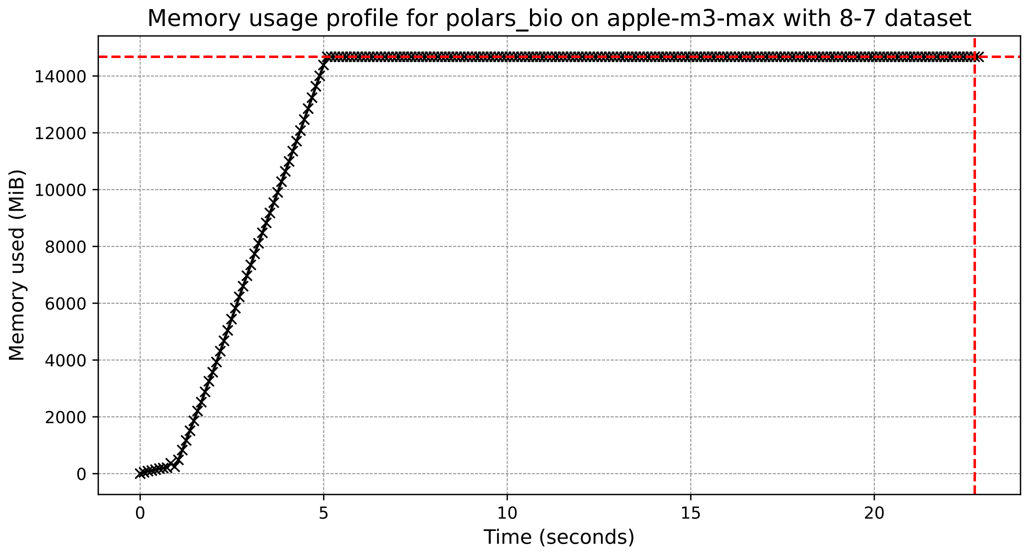
Memory usage profile for polars_bio on apple-m3-max with 8-7 dataset
Memory used (MiB)
Time (seconds)
0
2000
4000
6000
8000
10000
12000
14000
0
5
10
15
20

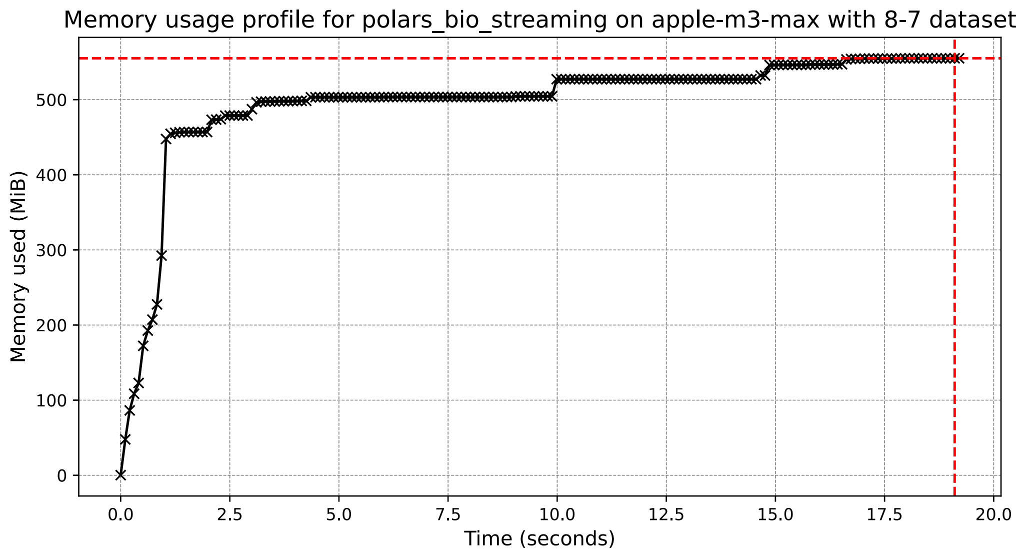

Memory usage profile for polars_bio_streaming on apple-m3-max with 8-7 dataset
Memory used (MiB)
Time (seconds)
0
100
200
300
400
500
0.0
2.5
5.0
7.5
10.0
12.5
15.0
17.5
20.0


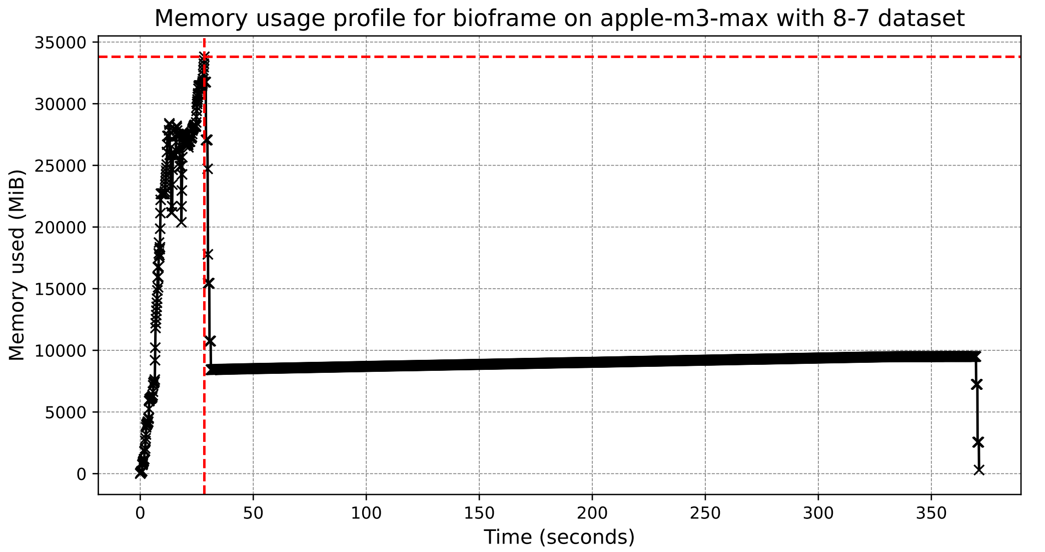

Memory usage profile for bioframe on apple-m3-max with 8-7 dataset
Memory used (MiB)
Time (seconds)
0
5000
10000
15000
20000
25000
30000
35000
0
50
100
150
200
250
300
350

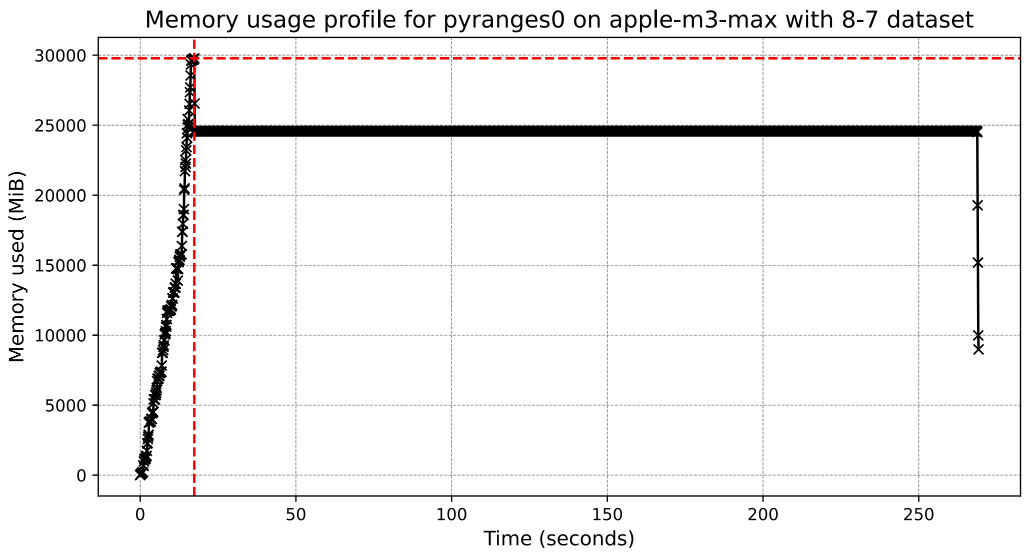

Memory usage profile for pyranges0 on apple-m3-max with 8-7 dataset
Memory used (MiB)
30000
25000
20000
15000
10000
5000
0
0
50
100
150
200
250
Time (seconds)


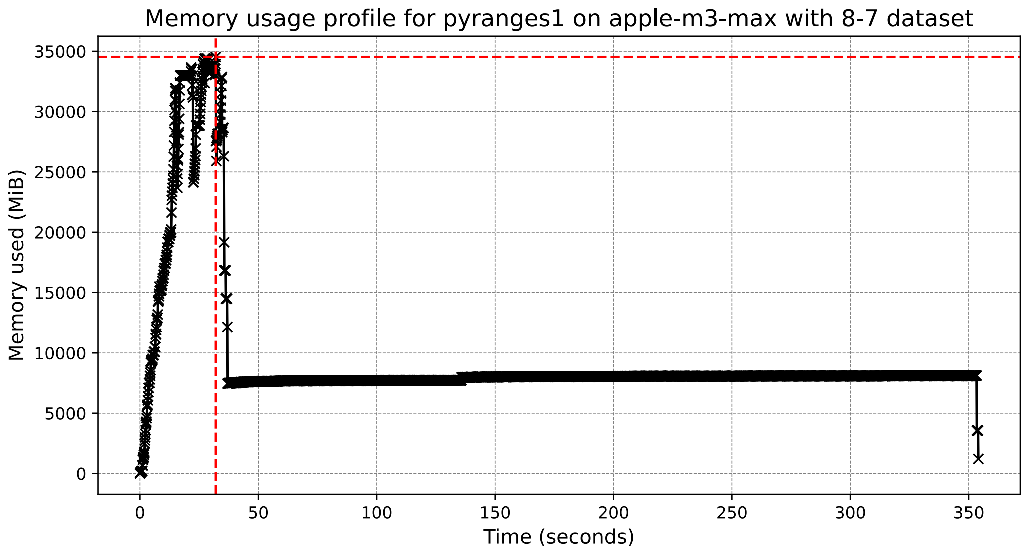

Memory usage profile for pyranges1 on apple-m3-max with 8-7 dataset
Memory used (MiB)
35000
30000
25000
20000
15000
10000
5000
0
0
50
100
150
200
250
300
350
Time (seconds)

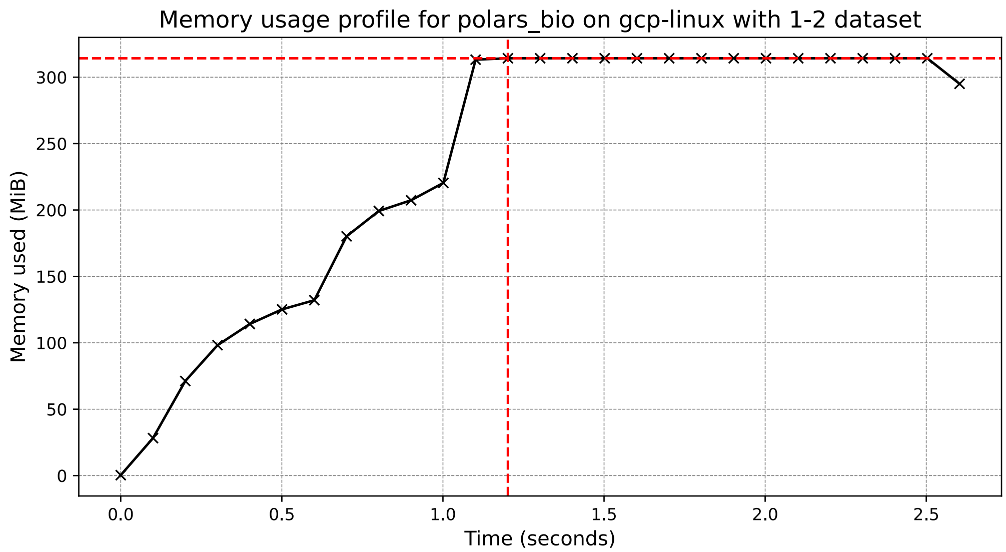
Memory usage profile for polars_bio on gcp-linux with 1-2 dataset
Memory used (MiB)
Time (seconds)
0
50
100
150
200
250
300
0.0
0.5
1.0
1.5
2.0
2.5

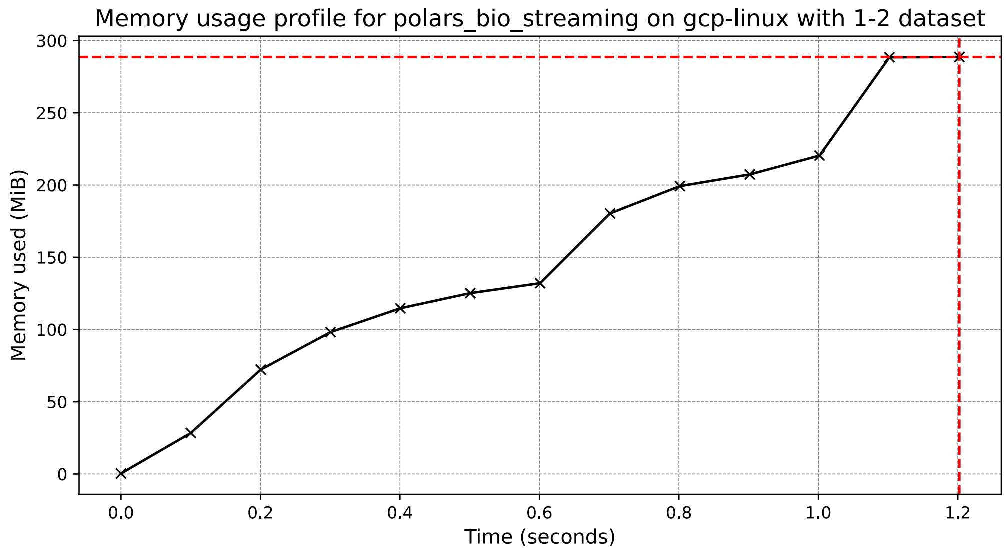
Memory usage profile for polars_bio_streaming on gcp-linux with 1-2 dataset
Memory used (MiB)
Time (seconds)
0
50
100
150
200
250
300
0.0
0.2
0.4
0.6
0.8
1.0
1.2

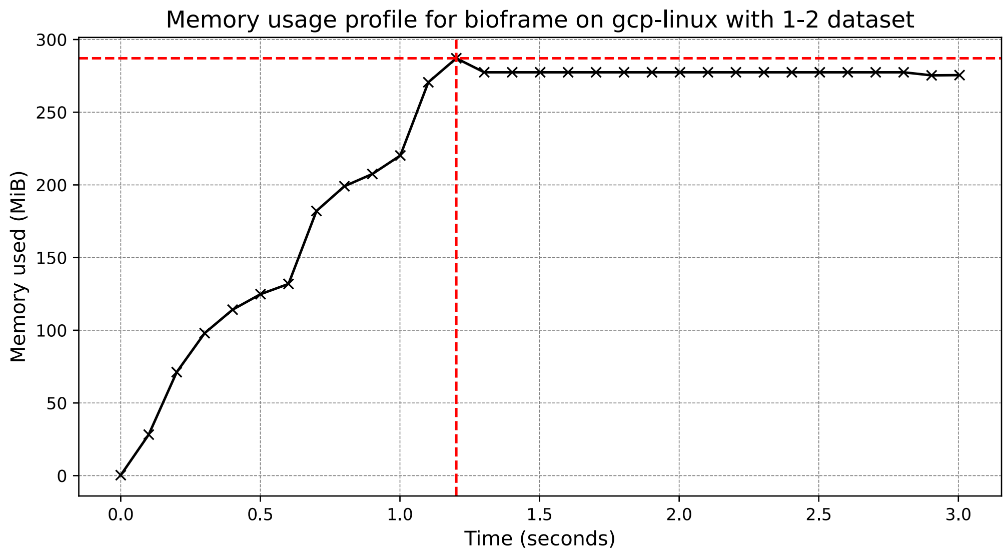
Memory usage profile for bioframe on gcp-linux with 1-2 dataset
Memory used (MiB)
Time (seconds)
0
50
100
150
200
250
300
0.0
0.5
1.0
1.5
2.0
2.5
3.0

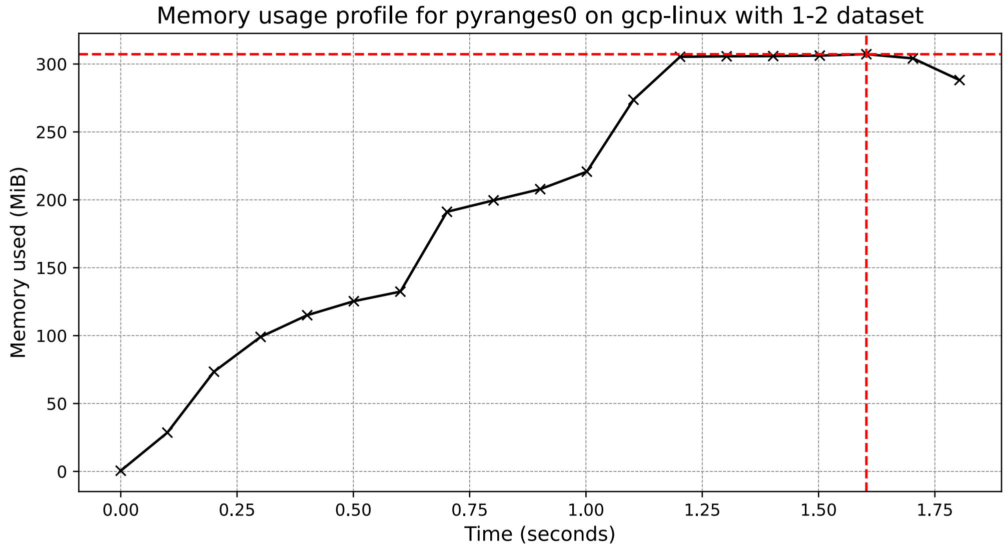
Memory usage profile for pyranges0 on gcp-linux with 1-2 dataset
Memory used (MiB)
Time (seconds)
0
50
100
150
200
250
300
0.00
0.25
0.50
0.75
1.00
1.25
1.50
1.75

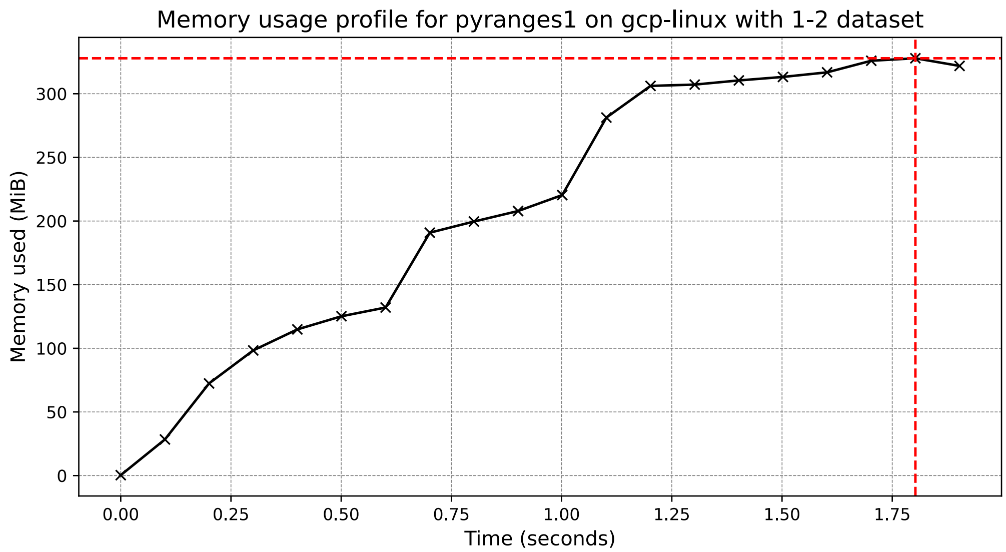
Memory usage profile for pyranges1 on gcp-linux with 1-2 dataset
Memory used (MiB)
Time (seconds)
0
50
100
150
200
250
300
0.00
0.25
0.50
0.75
1.00
1.25
1.50
1.75

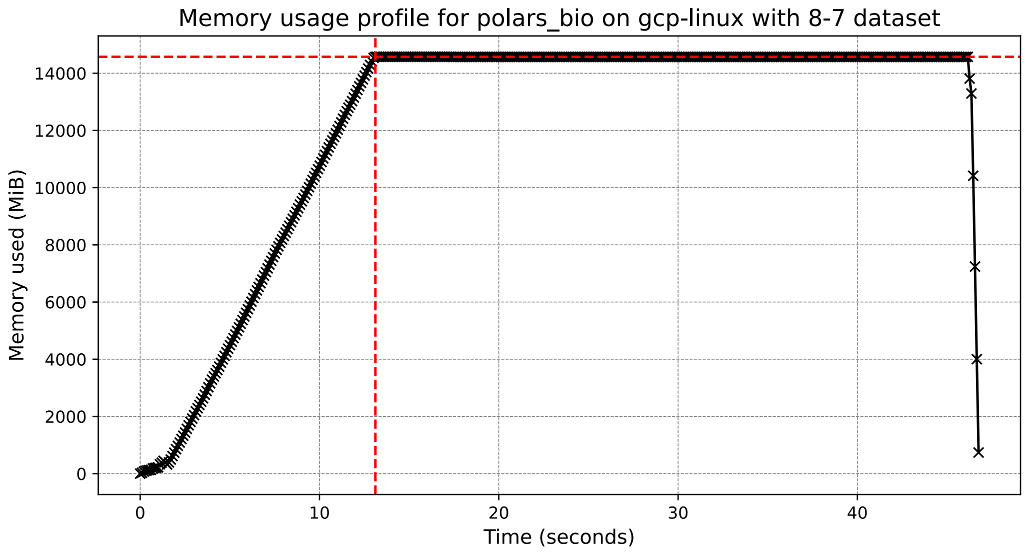
Memory usage profile for polars_bio on gcp-linux with 8-7 dataset
Memory used (MiB)
Time (seconds)
0
2000
4000
6000
8000
10000
12000
14000
0
10
20
30
40

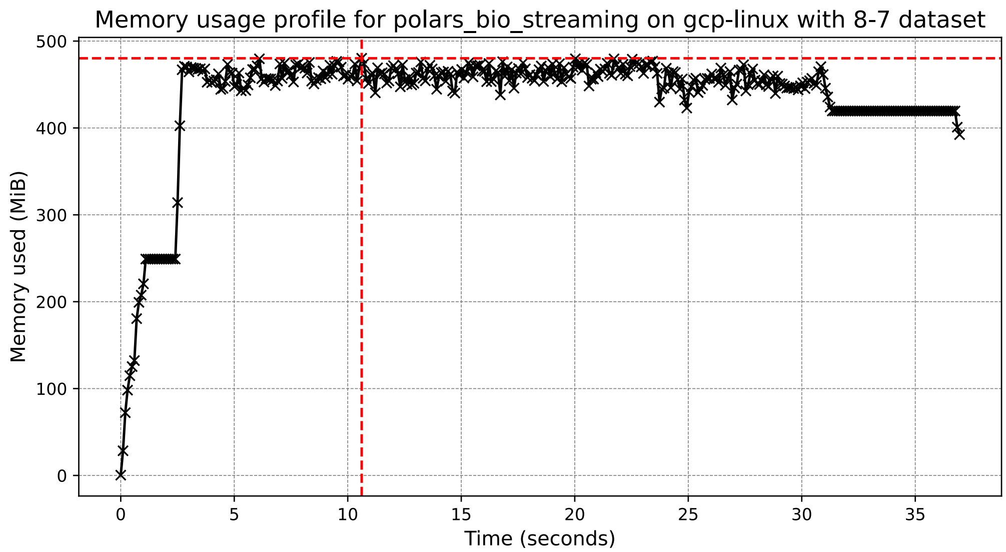

Memory usage profile for polars_bio_streaming on gcp-linux with 8-7 dataset
Memory used (MiB)
Time (seconds)
0
100
200
300
400
500
0
5
10
15
20
25
30
35


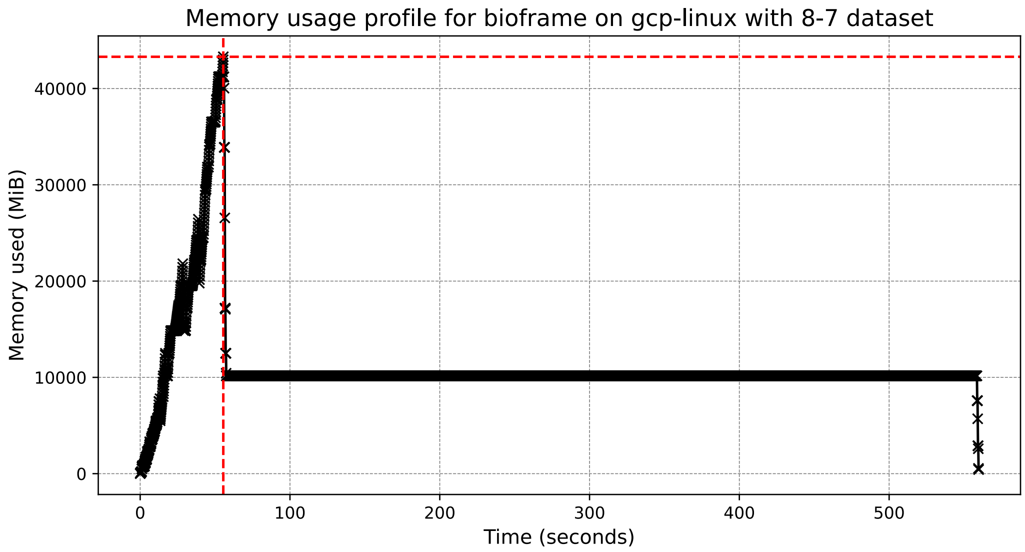

Memory usage profile for bioframe on gcp-linux with 8-7 dataset
Memory used (MiB)
Time (seconds)
0
10000
20000
30000
40000
0
100
200
300
400
500

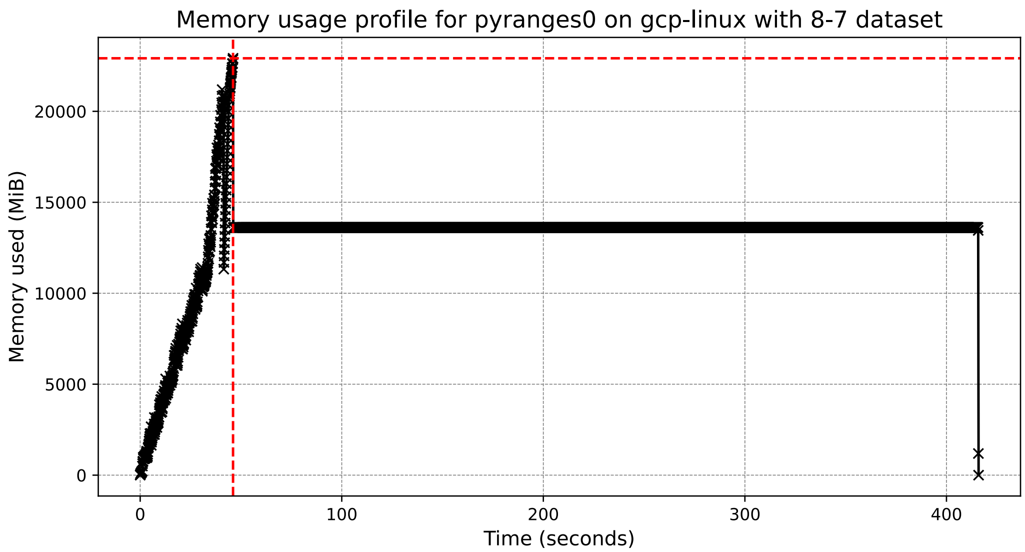
Memory usage profile for pyranges0 on gcp-linux with 8-7 dataset
Memory used (MiB)
0
5000
10000
15000
20000
Time (seconds)
0
100
200
300
400

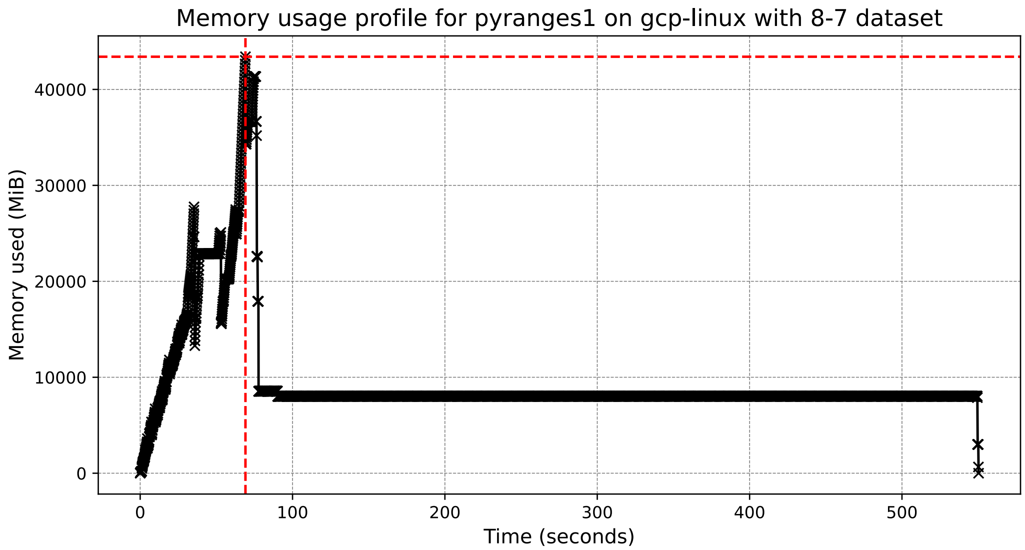
Memory usage profile for pyranges1 on gcp-linux with 8-7 dataset
Memory used (MiB)
0
10000
20000
30000
40000
Time (seconds)
0
100
200
300
400
500
